# Random sampling of ligand arrangements on a one-dimensional lattice

**DOI:** 10.1101/2024.02.17.580696

**Authors:** Dibyajyoti Mohanta, Albertas Dvirnas, Tobias Ambjörnsson

## Abstract

We introduce a transfer-matrix-based sequential sampling scheme for generating random samples of ligand arrangements on one-dimensional templates. The number of ligand types is arbitrary, the binding constants can have positional dependence, and cooperativity parameters are included. From the random arrangements, any (linear or non-linear) observable can be calculated using sample averaging. We provide a publically available software with a computational time that scales linearly with the lattice size. As an example application, we study the competitive binding of three ligand types (the sequence-specific binder netropsin, YOYO-1 and ethidium bromide) to a DNA molecule.

## I. INTRODUCTION

Ligand binding to long linear polymers like DNA plays important roles for biological function^1^. In processes such as gene regulation and DNA replication, ligands bind to DNA and linear proteins and subsequently induce changes in both chemical and mechanical properties, as well as affect the binding affinity of other ligands^2–4^. The site-specific binding of ligands, interactions between bound ligands (cooperativity), binding site (DNA) elongation per intercalated ligand molecule, etc., affect the binding mechanism and related biological functions of living systems^5–7^.

It has been a long digging field among physicists and chemists to study the binding of ligands to homogeneous lattices (the binding constants are the same for all sites), in contact with a bulk reservoir, using the framework of equilibrium statistical mechanics^8–11^. J. D. McGhee and P.H. von Hippel’s seminal theoretical work has served as an inspiration for subsequent studies in the field of ligand binding, which explores the concepts of specific and non-specific ligand binding to one-dimensional lattices^9^. The generating function approach offers an alternative method for predicting binding isotherms of multiple ligand binding to one-dimensional lattice^12–14^. The main focus has been on predicting binding isotherms, i.e., the average binding probability as a function of ligand bulk concentration, since this kind of observable can be measured using simple titration experiments.

For binding to heterogeneous lattices, the transfer matrix formalism (and variant thereof) emerges as a promising tool for investigating specific or non-specific binding, co-operativity, etc., on the ligand-lattice binding^15–17^. Heterogeneity can, for instance, occur in ligand binding to DNA molecules, due to ligand affinities being sensitive to DNA sequence. Recently, V.B. Teif and K. Rippe proposed a transfer-matrix-based approach for modelling the binding of different protein types to a one-dimensional lattice (DNA molecule)^15,16^. A. Nilsson and co-workers utilized the transfer matrix approach to study the competitive binding of multiple ligands to a heterogeneous one-dimensional lattice, which accurately reflects the underlying sequence signature of DNA^17^. In the latter study, the main focus was on calculating average binding probabilities along the DNA at a resolution set by the optical point spread function (PSF) of the imaging system. This kind of average probability (smeared by the PSF of a camera) can be experimentally measured in the field of competitive-binding (CB) optical DNA mapping (ODM), where fluorescent-stained DNA molecules are stretched using nanochannels and then imaged^17–19^. The CB method utilizes non-fluorescent netropsin and fluorescent YOYO-1 dyes to compete for binding sites on stretched DNA. YOYO-1 binds non-specifically, whereas the netropsin molecules exhibit high specificity for ATrich regions, out-competing YOYO-1 for AT sites. This results in a intensity variation along the DNA (AT/GC) that reflects the underlying sequence. Whereas the transfer matrix approach is effective at calculating average probabilities^17^, it remains a challenge to effectively compute full distributions (not just averages) of arbitrary observables of relevance in single-molecule studies.

A conceptually simple approach for computing arbitrary observables for ligand-binding systems is to generate random samples of ligand arrangements. While conceptually simple, naive approaches to this task, such as randomly positioning of ligands and then discarding unphysical arrangements (an arrangement where any two ligand overlapping, for instance) become computationally inhibitive for large systems (almost all arrangements would be discarded). Moreover, previous generating function based approaches deal with a single type of ligand and the computational time scale exponentially with lattice size^20^. Other combinatorial methods overlook important details in binding mechanisms like sequence specificity, and multi-ligand (of different sizes and kinds) binding.^10,21^.

In here, we introduce a new approach which is computationally fast (computational times scale linearly with the lattice size), where we treat ligand binding as a sequential process which utilizes the transfer matrices. Consequently, a random configuration of bound and free ligands is generated along the lattice. By averaging numerous such random configurations, we can effectively capture the precise theoretical probabilistic nature of the binding process, and at ease calculate any observable through simple sample averaging. To our knowledge, this study, for the first time carries out sequential sampling for generating random samples for multiple ligand binding to a heterogeneous lattice using explicit transfer matrices. We provide a MATLAB software (with a a graphical user interface, GUI), which generates random positional arrangements of ligands using our method. Our example application involve ligand binding to DNA, and therefore our software takes as input FASTA files with DNA sequences, along with a list of ligand bulk concentrations, sizes, binding constants and cooperativity parameters.

The modelling of ligand binding to one dimensional lattices is but one example of a statistical mechanics problem, which can be described using transfer matrices. In fact, any one dimensional statistical mechanics problem with short-range interactions can be treated using transfer matrices. As a consequence, our method for generating random samples can straightforwardly be extended beyond ligand binding.

## II. METHODS

The system at hand is illustrated in Fig. 1a). A onedimensional lattice of *N* sites is surrounded by a “bulk” consisting of *S* types of ligands at concentrations *c*^(*s*)^. We denote by *i* (*i* = 1, 2, …, *N*) the different lattice sites and use *s* (*s* = 1, 2, …, *S*) to denote different ligand types. A ligand of type *s* consists of *λ*^(*s*)^ monomers (i.e., when bound on the lattice, it occupies *λ*^(*s*)^ sites). The binding constant associated with binding of a ligand with its last monomer at site *i* (i.e., the ligand covers lattice sites [*i − λ*^(*s*)^ + 1, *i − λ*^(*s*)^ + 2, …, *i*]) for a ligand of type *s* is 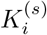. Following the enumeration scheme introduced by A. Nilsson, et al.^17^, we, without loss of generality, assign the statistical weight 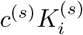 to the last monomer of the ligand, and a weight = 1 to the remaining bound monomers [one could, alternatively, but equivalently, assign weights, say 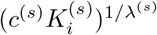 to each monomer, but this would make the transfer matrix below more complicated]. We also allow for cooperative binding between ligands of type *s* and of type *s*^*′*^ through the cooperativity parameters, 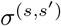. Double occupancy of lattice sites is not allowed. Note that we use subscript to denote sites and superscripts (with parenthesis) to denote ligand type (the use of superscript for increased clarity is different than the notation in Ref. 17). The problem at hand is now to generate random samples, i.e., random ligands arrangments on the lattice, for the system in Fig. 1a). These configurations must satisfy the constraint of no double site occupancy, and the associated sample average must yield equilibrium probabilities for a site to be occupied by a given ligand type.

**FIG. 1.**
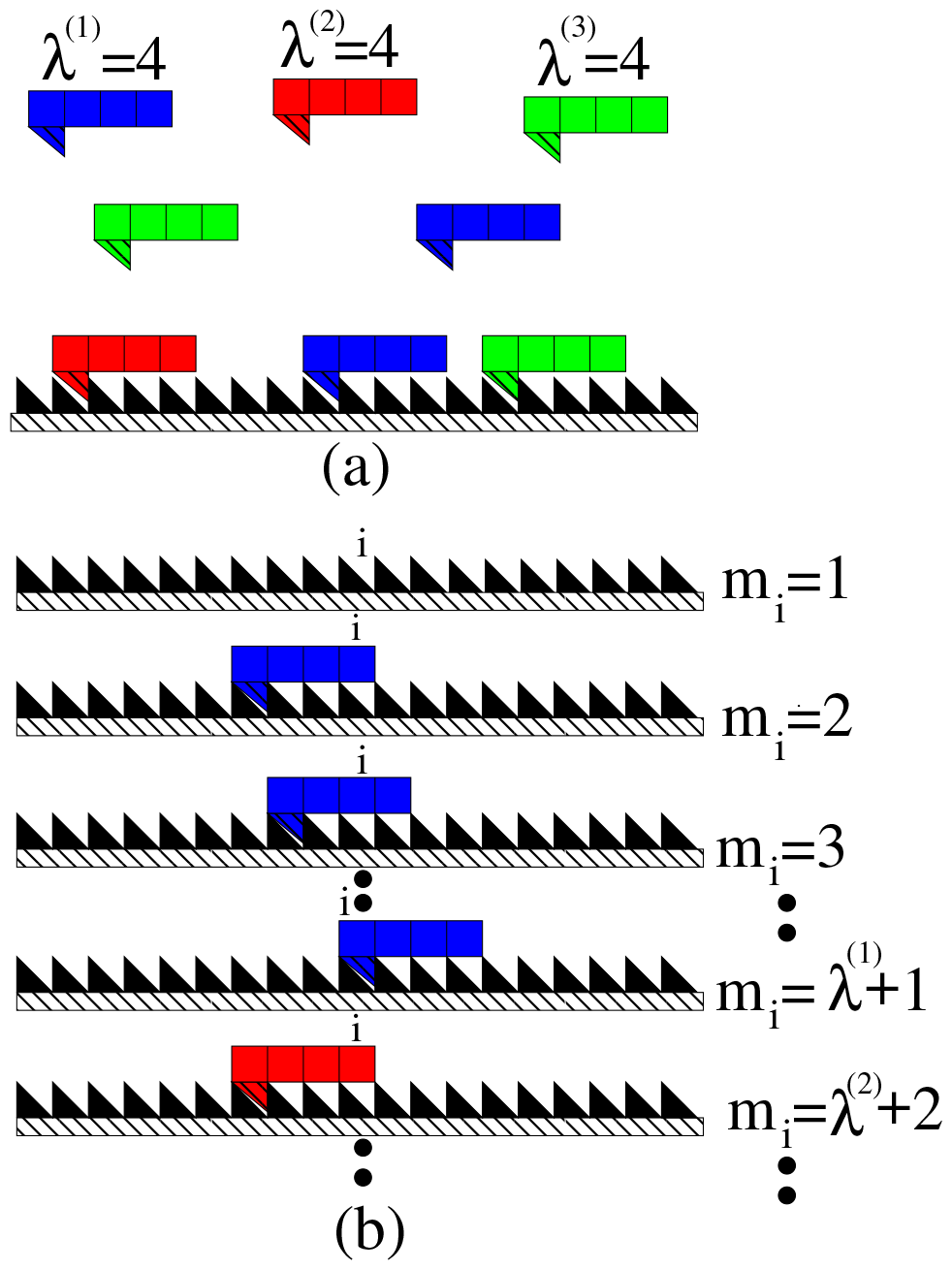
(a) Schematic illustration of the binding of different ligand types (here, three types) to a one dimensional lattice. No double occupancy of lattice site is allowed. (b) Schematic diagram of allowed states for a lattice site *i*. Each ligand consists of monomers, where a monomer covers one lattice site. Triangular indentations correspond to lattice sites. A ligand is bound when it fills up an indentation.

Our approach requires us to assign a state to each lattice site. Each site *i* can be in one of *M* possible states: the first state (*m*_*i*_ = 1) is that the site is empty, the second state (*m*_*i*_ = 2) is that the site *i* is occupied by the last monomer of a ligand of type 1, the third state (*m*_*i*_ = 3) is that the second-to-last monomer is occupying that site *i*, etc, see Fig. 1a) and Ref. 17). The total number of possible states for each lattice site is 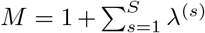. The output of our procedure is a state sequence

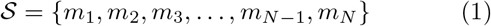

where the assigned states satisfy the constraints above.

The input to our method is a set of transfer matrices, **T**_*i*_. The matrix element *T*_*i*_(*m*_*i*_, *m*_*i*+1_) give the statistical weight for site *i* to be in state *m*_*i*_ provided site *i* + 1 is in state *m*_*i*+1_. In our case these are constructed, using the physical parameters (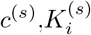 etc) as input, as described in detail in Ref. 17. As an example, for the case with the ligand types of sizes *λ*^(1)^ = *λ*^(2)^ = *λ*^(3)^ = 4, the transfer matrices become

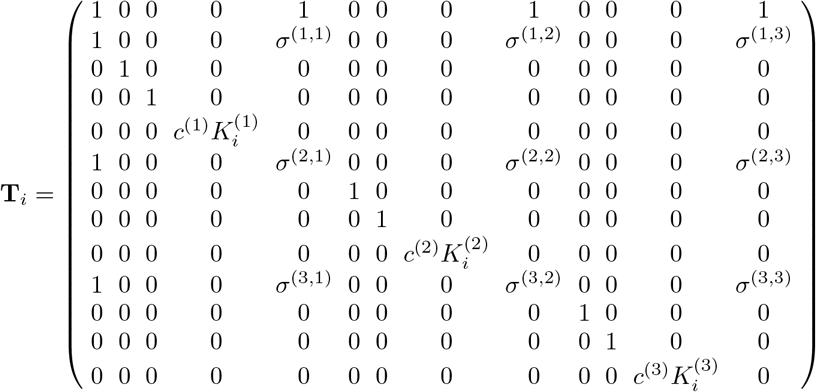

So, in this particular example we have 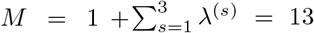 and majority of the elements of the *M × M* transfer matrix are zero. This follows from the fact that if a particular site *i* + 1 is occupied by last monomer of a ligand of type *s* = 1, then the next lattice site *i* cannot also be occupied by the last monomer of another type 1 ligand (if *λ*^(1)^ ≥ 2), etc.

The method is completed by also providing binary lists containing the allowed states for the last and first lattice sites^17^. Here, we simplify notation, and introduce two dummy sites (0 and *N* + 1) outside of the lattice to the left and right. We then require that these dummy sites are vacant, i.e.,

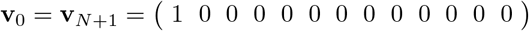

This procedure is equivalent to the procedure in Ref. 17, where only a dummy site to the right was used.

Our general procedure for generating random state sequences using a set of transfer matrices as input is described in appendix B. Our method uses the probability, *p*(*m*_*i*_ | *M*_prev,*i*_), of the current site *i* to be in the state *m*_*i*_ given all “previous” sites were in states: *M*_prev,*i*_ = {*m*_*i*+1_, …, *m*_*N*_}. We then draw a random state for site *i* based on *p*(*m*_*i*_ | *M*_prev,*i*_), where we start at site *N* and continue until we reach site 1. The quantity *p*(*m*_*i*_ | *M*_prev,*i*_) is defined in Eq. A4, but is in practice calculated recursively using Eqs. A5 and A6. These recursions involve two sets vectors **u**^*L*^(*i*), and **u**^*R*^(*i*) (*i* = 1, …, *N* + 1) and associated normalization constants *n*^*L*^(*i*) and *n*^*R*^(*i*) (needed to avoid numerical problems for large *N*). Note that the method in the appendix B works for arbitrary choices of the transfer matrices (i.e., arbitrary statistical mechanics problem with short-range interactions).

We here present a computationally faster version, the “fast-forward” approach, of the general method, which utilizes the fact that most of the elements in the transfer matrices are zero, i.e., in certain cases, the next state in the state sequence is “immediately known” without computations. Our method is targeted towards ligandbinding problems where double-occupancy of sites is forbidden. For instance, if we are in state *m*_*i*_ = *λ*^(1)^ + 1 = 5 (the site is occupied by the last monomer of ligand 1), there is no need to calculate the probability from Eq.A5, as the next site (*i* − 1) must be in state *m*_*i−*1_ = 4 with probability 1 (see Appendix A). Subsequently, the next two sites, *i −* 2 and *i −* 3, will be in states *m*_*i−*2_ = 3 and *m*_*i−*3_ = 2 (for *λ*^(1)^ = 4). It should be mentioned here that this ‘fast-forward’ approach is a special case of the general approach described in Appendix B and does not work for arbitrary choices of the transfer matrix T, as the general approach does.

Our fast-forward algorithm starts by initiating the vectors **u**^*L*^(0) and **u**^*R*^(*N* + 1) as **u**^*L*^(0) = **v**_0_*/*|**v**_0_| and **u**^*R*^(*N* +1) = **v**_*N*+1_*/*|**v**_*N*+1_|. We also initialize site counter *i* = *N*.

Our method is then:

1. Calculate the “left” vectors and normalization constants for all sites. Begin at 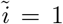 and calculate 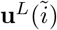 and 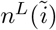 based on Eq. A6 until 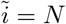.
2. Calculate the quantities **u**^*R*^(*i*) and *n*^*R*^(*i*) in Eq. A6 using the “previous” values (from step *i* + 1) as input. From these, the conditional probability *p*(*m*_*i*_|*M*_prev,*i*_) is obtained through Eq. A5.
3. If the calculated probability *p*(*m*_*i*_|*M*_prev,*i*_) yields more than one non-zero value for being one of the possible states, pick a random state, *m*_*i*_ with probabilities given by *p*(*m*_*i*_|*M*_prev,*i*_). To this end, we generate a uniformly distributed random number *r* in the range [0,1] and perform a binary search in the cumulative sum of probabilities^25^.
4. We then distinguish between two cases:
  - If site *i* is chosen to be in any states defined by the last monomer of any ligand type, we apply the “fast-forward” approach. In this scenario, once a site is occupied by the last monomer of a given ligand of type *s*, the subsequent sites will be occupied by the second last to the first monomers of that particular ligand. For each of these “fast-forwarding” steps **u**^*R*^(*i*) is equal to the state vector 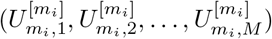 (where site *i* + 1 is in state *m*_*i*+1_ = *m*_*i*_ + 1; see Appendix A). Update the site counter *i* accordingly and return to step 2.
  - If the chosen state is one where there is no ligand at site *i* (*m*_*i*_ = 1), then update the site counter *i → i −* 1 and return to step 2.

We iterate through the above steps until we reach *i* = 1. The algorithm above allows us to generate random states along the lattice sites for a single run. Once the complete state sequence is obtained, we precisely determine whether a certain ligand type (*s*) occupies a specific site *i* (a binary outcome = 1) in the lattice or not (a binary outcome = 0). From many (*n*_*run*_) random realizations any observable of interest can be calculated using sample averages.

In our Results, we focus on illustrating the result of our algorithm for the site occupancy probability of different ligand types. To calculate this observable, we calculate sample average, and associated standard deviations, over all outcomes (0s and 1s) from the *n*_*run*_s for each site *i* for ligands of type *s*. For validation, we compare our average probabilities of ligand binding to the exact site occupancy probability calculated using the procedure described in Ref. 17.

## III. RESULTS

We now illustrate our method using *λ*-phage DNA molecule of size 48,502 bp with 55% AT content as “lattice” template for ligand binding^26^. As ligands, we use 4-base pair (lattice unit) length Netropsin and YOYO-1 with concentration ratio relevant to the experiment as ligand 1 (*s* = 1) and ligand 2 (*s* = 2) with 6 and 0.02 *μM* respectively^17^. As ligand 3 we have included four base-pair long Ethidium bromide with a constant binding constant 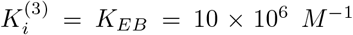 and concentration of 0.02 *μM*. The DNA concentration in our calculation is 0.2 *μM*. Here, YOYO-1 (ligand 2) exhibited a constant binding constant 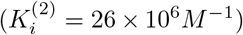, while Netropsin’s binding involved a sequence-specific set of 4^4^ = 256 binding constants 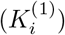 extracted originally from percentage fluorescence intensity chart from the article by D. L. Boger, et. al.^27^ and then calculated in supplementary information of article^28^. The cooperativity parameters 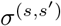 (*s, s*^*′*^ = 1, 2, 3) are set to 1. The parameter values are summarized in Table I.

**TABLE I.**
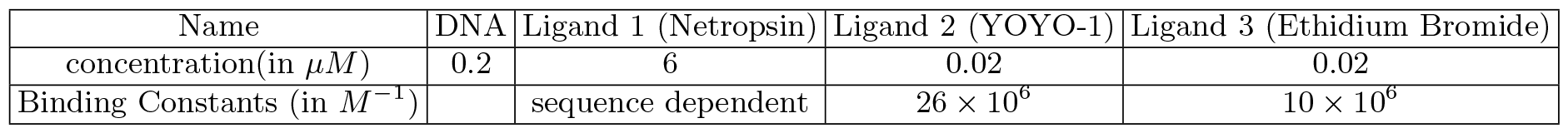
Ligand parameters as used in the Results section. The nine cooperativity parameters are set as *σ*^(*i,j*)^ = 1 for *i, j* = 1, 2, 3 within the matrix **T**_**i**_.

Figure 2 (2) displays positional maps of ligands (s=1,2,3) from 10 different stochastic simulations along the DNA sequence for the first 300 base-pairs stacked vertically. The figure indicates that, due to the high concentration and binding constant of ligand 1 (Netropsin), the majority of sites are occupied by it (shown in blue). Ligand 2 (YOYO-1), with a higher binding constant (Table I.) than ligand 3, is the second most prevalent, represented by the red colour. Ligand 3 (Ethidium bromides) is observed the least populated in the maps (green). It is worth noting that the lattice template is not fully crowded by ligands, as occasionally free sites in between ligands are depicted by the black colour.

**FIG. 2.**
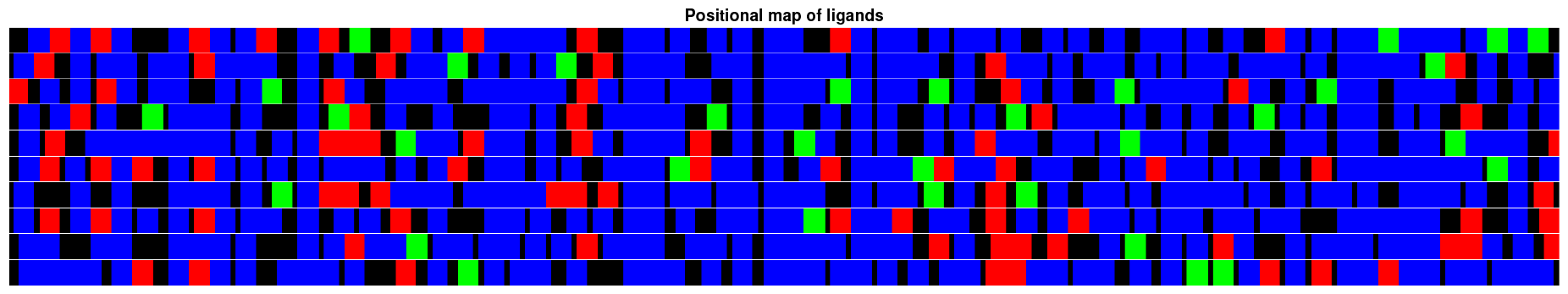
Positional map of ligands comprising of 10 random state sequences along the DNA (from 1 to 300bp) are stacked together (along vertical axis), where the black colour bar shows an empty site, the blue bar represents Netropsin (ligand 1), the red represents represents YOYO-1 molecule (ligand 2) and the green represents ethidium bromide (ligand 3).

To validate our method, we compare the average of the random state sequences from our method for 1000 *n*_*run*_s to the exact predicted mean binding probabilities, see Ref. 17. From the figure 3, we can see that the mean of the stochastic simulation (red dashed) shows excellent agreement with the exact probabilistic binding (blue solid line). Note, however, that from our stochastic simulation, we can not only calculate average probabilities, but also standard deviations and any other “non-linear” observables.

**FIG. 3.**
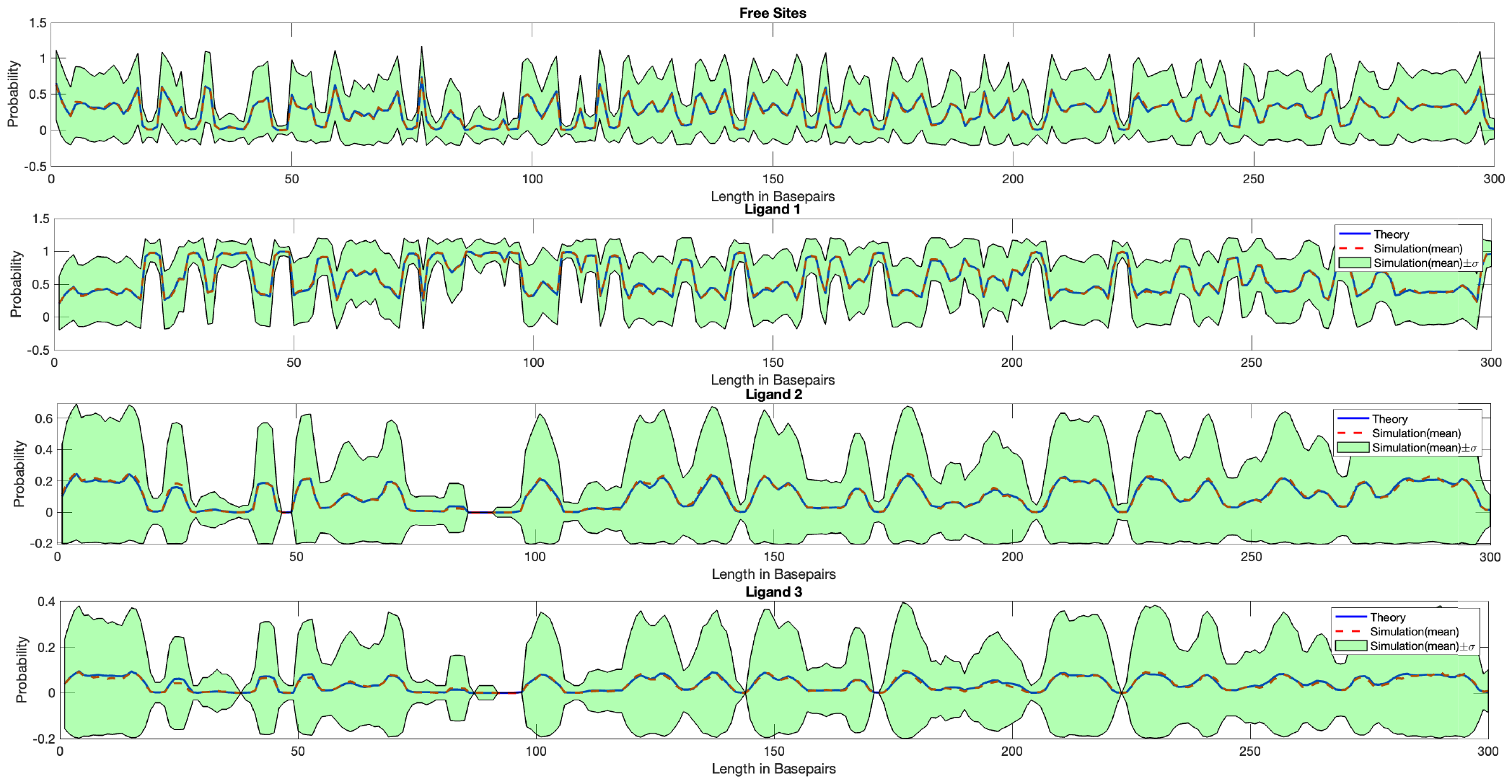
Comparisons of normalized stochastic binding to theoretical exact binding results for (a) Free sites, (b) Netropsin (ligand-1), (c) YOYO-1 (ligand 2), and (d) ligand 3, along the DNA length (0 → 300 bp). The standard deviation (*σ*) depicted in the figure is computed from 1000 randomly generated samples for each ligand type across the lattice.

The computational time for one random realization for the full lambda-phage DNA was less than a second on a laptop computer with an Apple M1 chip with 16 GB RAM memory. Because of the nature our algorithm (sequential sampling), the computational time scales linearly with the lattice size, *N*. The software and code (GUI version and non GUI version) supporting the finding of this article are available at https://github.com/dibyajyoti41/ligandbinding

## CONCLUSION

We have introduced a computationally fast method for generating random samples of ligand arrangements on one-dimensional lattices. Once this random configurations are averaged over the total runs, we retrieve the equilibrium probability of ligand binding from exact calculation. Our approach of generating random state sequences is both rapid and precise, making it applicable to various ligands and protein binding mechanisms on one-dimensional homo or hetero sequence templates. Our MATLAB-based software comes both as in a “script” format and with a GUI. The GUI is user-friendly, guiding users through an interactive dialogue box to input necessary parameters during the simulation and generates outputs of binding probabilities for easy interpretation. We hope this study, along with the accompanying software, will be valuable for researchers, enabling fast and precise calculations of ligand binding to any DNA, linear protein or other one-dimensional substrates.

## Appendix A The conditional probability

We here provide an explicit expression for conditional probability for lattice site *i* to be in state *m*_*i*_ given that the site *i* + 1, … *N* were in states *M*_prev,*i*_ = *{m*_*i*+1_, … *m*_*N*_ *}*.

For each site we introduce an *M* × *M* transfer matrix **T**_*i*_ with elements *T*_*i*_(*m*_*i*_, *m*_*i*+1_)^15,16^, which give the statistical weight for site *i* to be in state *m*_*i*_ provided site *i* + 1 is in state *m*_*i*+1_. In this way, we have that the statistical weight of a particular state sequence, 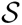, is:

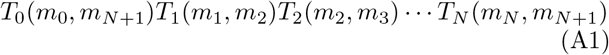

where matrices *T*_0_(*m*_0_, *m*_*N*+1_) and *T*_*N*_ (*m*_*N*_, *m*_*N*+1_) includes the two dummy/vacant states *m*_0_ and *m*_*N*+1_ (*m*_0_ = *m*_*N*+1_ = 1) at sites 0 and *N* + 1.

Our simulation procedure makes use of the conditional probability, *P* (*m*_*i*_|*M*_prev,*i*_) that the “next” site, *i*, is in state *m*_*i*_, given that all “previous” sites were in states *M*_prev,*i*_ = {*m*_*i*+1_, *m*_*i*+2_, …, *m*_*N*_*}*. In terms of the transfer matrices, we have

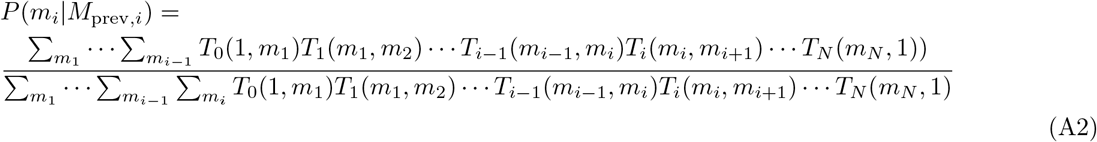

Note that we keep (*m*_*i*+1_, *m*_*i*+2_, …, *m*_*N*_, *m*_*N*+1_) “fixed”, i.e., those variables are not summed over. Also, note that we have the normalization condition 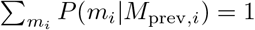 satisfied, as it should.

To write the equation above in matrix form, we introduce the *M* projection matrices defined as (*q* = 1, … *M*)

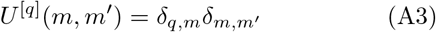

Here, *δ*_*n,n*_*′* is the Kronecker delta. We here use square bracket in the superscript to avoid confusion with the ligand labels (see main text).

As an example, for *S* = 3 (three ligand types) and *λ*_1_ = *λ*_2_ = *λ*_3_ = 4 as in the Results section of the main text, we have

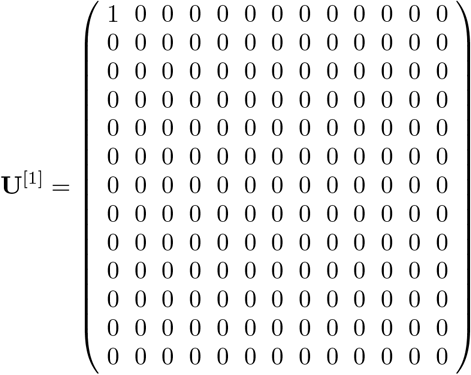

Similarly for **U**^[2]^, **U**^[3]^, and so on.

Using Eq. (A3) and using the dummy state vectors **v**_0_ and **v**_*N*+1_, we can write Eq. (A2) according to:

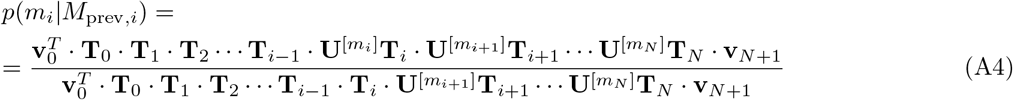

To avoid numerical issues associated with repeated matrix multiplications, we adopt the normalization procedure in^17^ for evaluating *p*(*m*_*i*_ |*M*_prev,*i*_) numerically. To this end, we rewrite Eq. (A4) on the form:

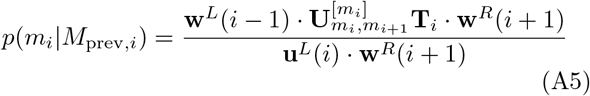

where we have the recursion relations:

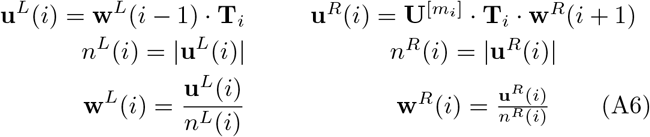

The set of equations above constitutes recursion relations for *p*(*m*_*i*_|*M*_prev,*i*_), which can be evaluated numerically, using ‘initial’ conditions, 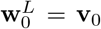 and 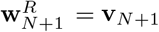.

## Appendix B General approach for generating random samples

We introduce a general method for generating state sequences on the form in Eq. (1). Our method takes as input a number of transfer matrices and binary vectors listing allowed states for the ends, and then generates several random state sequences 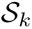 (*k* = 1, …, *n*_*runs*_). The method makes use of *p*(*m*_*i*_ |*M*_prev,*i*_) (see previous appendix), which is evaluated recursively.

We initiate the vectors **u**^*L*^(0) and **w**^*R*^(*N* + 1) at the two ends (*i* = 0 and *i* = *N* + 1). We set the counter at *i* = *N*.

Our procedure is then as follows :

1. Calculate “left” vectors and normalization constants for all sites. Begin at 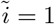. Calculate 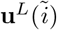 and 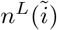 from 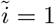 to 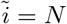 based on the recursive relation, Eq. A6.
2. Calculate **w**^*R*^(*i*) and *n*^*R*^(*i*) in A6 using the “previous” (*i* + 1) values as input. Then calculate the conditional probability *p*(*m*_*i*_|*M*_prev,*i*_), where *M*_prev,*i*_ = *{m*_*i*+1_, …, *m*_*N*_ *}* using Eqs. A5 and A6.
3. Pick a random state, *m*_*i*_ with probabilities given by *p*(*m*_*i*_ |*M*_prev,*i*_). In practice, we generate a uniformly distributed random number *r* in the range [0,1] and a binary search is used in the cumulative sum of probabilities to determine the state at the current site *i*^25^.
4. Update *i → i −* 1 and return to step 2.

We repeat steps 2-4 above until *i* = 1.

The procedure above yields our first (*k* = 1) random state sequence and is then repeated *n*_*runs*_ times. From these random state sequences we can then calculate sample averages and compare the results with if available the exact calculated probability of states along the sequence.

## Notes

### Competing Interest Statement

The authors have declared no competing interest.

